# Mechanism of evolution by genetic assimilation: Equivalence and independence of genetic mutation and epigenetic modulation in phenotypic expression

**DOI:** 10.1101/242206

**Authors:** Ken Nishikawa, Akira R. Kinjo

## Abstract

Conrad H. Waddington discovered the phenomenon of genetic assimilation through a series of experiments on fruit flies. In those experiments, artificially exerted environmental stress induced plastic phenotypic changes in the fruit flies, but after some generations the same phenotypic variant started to appear without the environmental stress. Both the initial state (where the phenotypic changes were environmentally induced and plastic) and the final state (where the phenotypic changes were genetically fixed and constitutive) are experimental facts. However, it remains unclear how the environmentally induced phenotypic change in the first generation becomes genetically fixed in the central process of genetic assimilation itself. We have argued that the key to understanding the mechanism of genetic assimilation lies in epigenetics, and proposed the “cooperative model” in which the evolutionary process depends on both genetic and epigenetic factors. Evolutionary simulations based on the cooperative model reproduced the process of genetic assimilation. Detailed analysis of the trajectories has revealed genetic assimilation is a process in which epigenetically induced phenotypic changes are incrementally and statistically replaced with multiple minor genetic mutations through natural selection. In this scenario, epigenetic and genetic changes may be considered as mutually independent but equivalent in terms of their effects on phenotypic changes. This finding rejects the common (and confused) hypothesis that epigenetically induced phenotypic changes depend on genetic mutations. Furthermore, we argue that transgenerational epigenetic inheritance is not required for evolution by genetic assimilation.

## 1 Introduction: Phenotype-driven evolution

In the 1950s, Waddington discovered the phenomenon of genetic assimilation (Waddington 1957). Some environmental stress artificially exerted on fruit flies during a certain developmental stage induces plastic phenotypic changes in the adult flies. By artificially selecting these phenotypic variants over some generations, the fruit flies eventually come to express the mutant phenotype even without the environmental stress. That is, the mutant phenotype is genetically fixed or assimilated (Scharloo 1991). We can observe the following characteristics in genetic assimilation:

a. A strong environmental stress induces phenotypic changes in multiple individuals of a population.
b. The mutant phenotype in the first generation becomes, after a series of artificial selection over generations, genetically fixed (genetic assimilation).
c. Genetic assimilation completes in a relatively small number of generations. In other words, genetic assimilation is a rapid evolutionary process.

Waddington’s experiments were carried out for at least two types of mutant phenotypes, *cross-veinless* wing (Waddington 1953) and *bithorax* (Waddington 1956), both of which demonstrated genetic assimilation. Furthermore, they were performed based on a then-well-known phenomenon called “phenocopy” (Goldschmidt 1938) which is a phenomenon that external stimuli during ontogenesis induce phenotypic changes in the same generation that are indistinguishable from known genetic mutants. The initial phenotypic change is therefore considered as an acquired trait caused by the environmental stress (the point (a) above). But the change is somehow genetically fixed in the final stage. Waddington himself considered that genetic assimilation was caused by “developmental plasticity” (in today’s term), and explained it, rather metaphorically, as a change in the pathways of developmental process (which he called “canalization”) due to the environmental change.

The standard evolutionary theory, namely the Evolutionary Synthesis, assumes individuals with an advantageous *heritable* trait must preexist in the population in the initial stage of evolution, and the genetic mutations responsible for the trait are gradually fixed in the population over generational changes. As such, genetic assimilation (where the initial phenotypic change is not heritable) poses a serious difficulty on today’s evolutionary theories, and it is often avoided as a “rare” phenomenon (Laland et al 2014).

Among several alternative theories to the Evolutionary Synthesis, collectively called the “Extended Evolutionary Synthesis” (Pigliucci and Müller 2010), a major one is Evo-Devo (Müller 2007) which emphasizes the leading role of development in evolution. Evo-Devo advocates a decisive role of phenotype rather than genotype in evolution. This view is well phrased by West-Eberhard: “Environmental induction is a major initiator of adaptive evolutionary change. The origin and evolution of adaptive novelty do not await mutation; on the contrary, genes are followers, not leaders, in evolution.” (West-Eberhard 2003). The Evo-Devo view of evolution as represented by West-Eberhard is called “phenotype-driven evolution” in the present review.

West-Eberhard proposed “genetic accommodation” as a mechanism of phenotype-driven evolution (West-Eberhard 2003), which generalizes Waddington’s notion of genetic assimilation. Her monograph (West-Eberhard 2003) is full of circumstantial evidence supporting evolution by genetic accommodation. However, rather unfortunately, she could not explain explicitly how a (non-heritable) phenotypic change is replaced with genetic mutations as generations change (the point (b) above).

In order to clarify the problem, let us define “evolution” as follows:

> *Evolution is a process in which individuals whose phenotypes are adaptive to a given environment are naturally selected, and thereby fixing in the population the genotype that induces the adaptive phenotype.*

In this sense, genetic assimilation is clearly an example of evolution. The Evolutionary Synthesis cannot fully explain the phenomenon because the initial phenotypic change is not due to genetic mutations. On the other hand, West-Eberhard in particular, and the Extended Evolutionary Synthesis in general, do not explain how the initial non-genetic phenotypic change can be genetically fixed in the final stage.

If we reexamine West-Eberhard’s thesis in modern terms, we notice that this is a problem of epigenetics which tries to connect phenotypes with genotypes in molecular terms. It was before the dawn of modern epigenetics when West-Eberhard published her monograph in 2003. Thus, she was not in a historical position to enjoy the fruitful and vast results of epigenetics; but now we are. It is therefore our responsibility to overcome the problem West-Eberhard challenged but failed, and to review the evidence underpinning the evolutionary mechanism of genetic accommodation (we will focus on genetic assimilation henceforth) in molecular terms.

In the following, we first review basic notions of epigenetics and its role in evolution as well as some attempts to solve the aforementioned problem. However, we do not discuss transgenerational epigenetic inheritance (TGEI) which often comes into focus when epigenetics is related to evolution (Jablonka and Raz 2009): genetic assimilation can be explained without assuming TGEI.

## 2 Genome vs. Epigenome

The conventional gene-centric biology cannot properly treat the relationship between phenotype and genotype. The simple scheme that a gene determines a trait, or the one-to-one correspondence between genes and traits, has been rejected. Monozygotic twins, sharing the identical genomes but exhibiting notable phenotypic differences, suffice as a simple counterexample against the gene-centric view. On the other hand, epigenetics can consistently explain phenotypic differences resulting from the identical genomes through ontogenesis (Fraga et al 2005). Such phenomena stand out even more at the cellular level than at the individual level. That is, all the cells in an individual originate from a single fertilized egg and thus share the identical genome, nevertheless, their phenotypes are extremely diverse depending on their cell types, tissues and organs. According to epigenetics, these differences are due to the differences in their “epigenomes.”

The substance of epigenome is a set of chromatin modifications consisting of DNA methylations and histone modifications, which modulate the genome-wide regulation of gene expression. The chromatin modifications are also called epigenetic marks; these marks are “written” on the chromatin (addition of modifications), “read out” (recognition by chromatin binding proteins) or “deleted” (erasure of modifications). Epigenetic marks are persistent so that they are not easily deleted. In particular, the epigenetic marks of a mother cell are copied to its daughter cells through mitotic cell division. On the contrary, fertilized eggs go through a massive erasure of epigenetic marks called reprogramming (Reik et al 2001), and ontogenesis starts from the “null state” without epigenetic marks, although some part of genomic DNA are not reprogrammed in mammals and plants (genome imprinting) (Feil and Berger 2007; Ferguson-Smith 2011).

The epigenome, the information of which may be written, maintained, read out or deleted, may be considered as “meta information” (Ohta 2013) that is of a different kind than the genome itself. This situation may be schematically described as a double-layered structure in which the epigenome as meta information covers the genome as (primary) information (Fig. 1 in Nishikawa and Kinjo (2017)). While the epigenome regulates gene expression according to epigenetic marks (action of epigenome on genome; Nishikawa and Kinjo (2017)), the genome encodes chromatin modification enzymes and (non-coding) RNAs that construct and modify the epigenome (action of genome on epigenome). The latter action is clearly demonstrated in the process of cell differentiation where *de novo* chromatin modifications are incrementally added, starting from the stem cell to differentiated cells to establish cell type-specific epigenetic marks.

**Fig. 1.**
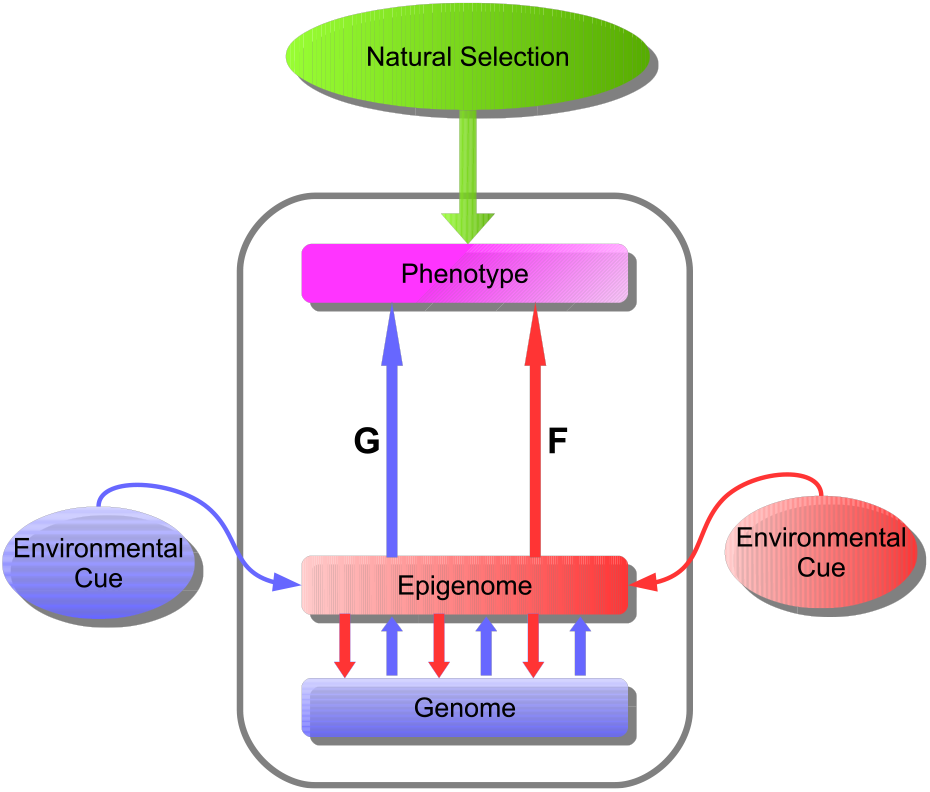
Schematic relationships among genome, epigenome and phenotype in a cell. Gene expression of the genome is regulated by the epigenome (chromatin modifications such as DNA methylation and histone marks), as indicated with downward red arrows towards the genome. The epigenome of a particular cell is specified by environmental effect as well as the genome which encodes chromatin modifying enzymes and non-coding RNAs, as indicated with upward blue arrows out of the genome. The phenotype of the cell arising both from genetic (G) and epigenetic (F) pathways is targeted by the natural selection. Pathway G including an anticipated environmental cue (blue arrow from the left) is genetically programmed so that its alteration needs the change in the genome (i.e., mutation in DNA sequence), while epigenetic pathway F arises from the epigenome that is affected with an unanticipated environmental cue (red arrow from the right) but without alternation in the DNA sequence.

An important feature of epigenome that contrasts with genome is that it can be altered by the influence of environment. An environmental change can alter the epigenome, which alters gene expression patterns, which in turn may trigger, for example, diseases. Such environment-induced epigenetic changes may also contribute to evolution, which we discuss next.

## 3 Two pathways to phenotype

Behind the macroscopic phenomena such as phenotypes, phenotypic varieties and phenocopy lie microscopic and complex epigenetic phenomena. The relationship between macroscopic phenotype and environment, and microscopic genome and epigenome can be schematically represented as in Fig. 1.

As already pointed out above, genome and epigenome comprise a double-layered structure from which two pathways lead to the phenotype. The pathway from genome through epigenome to phenotype (blue arrows in Fig. 1) represents a normal ontogenesis through cell differentiation. In this process, the epigenomes of cells are altered so that differentially expressed genes determine the differential phenotypes of the cells. Nevertheless, the genomic information (DNA sequence) itself remains unaltered (we do not consider mutagenesis in this paper).

Not all the phenotypes brought about by normal developmental processes are determined by the genome; some are modulated by epigenome depending on environmental conditions. A well-known example is the caste polymorphism of the honey bee where the intake of royal jelly during the larva period, which affects the epigenome, determines whether the adult bee becomes a queen or worker (Kucharski et al 2008; Weiner and Toth 2012). In this case, the branched developmental processes determined by the presence or absence of the environmental factor must have been evolutionarily acquired. The same argument can be applied to the blooming process of plants where it is necessary for the plants to experience cold winter before becoming ready to bloom (Ietswaart et al 2012; Jones and Sung 2014). There are many other examples of drastic morphological changes modulated by epigenetic mechanisms in response to environmental changes (Gilbert and Epel 2009; Pfennig et al 2010; Simon et al 2011). Although drastically affected by the environment, these examples are yet genetically programmed. The blue pathway to the phenotype in Fig. 1, which may or may not be epigenetically modulated, is called the “G pathway” as it is primarily genetically determined.

The other, red, pathway in Fig. 1 corresponds to *ad hoc* and phenotypically plastic responses to environmental stimuli. The origin of this pathway is the environment which leads through the epigenome to the phenotype. In contrast to the G pathway, the red pathway is not genetically programmed, but *ad hoc.* A phenotypic change due to the F pathway is limited to each individual and is not inherited to the next generation. In this sense, it is non-genetic, or literally epigenetic. In the following we call this pathway the “F pathway”.

Individuals’ responses by phenotypic plasticity may be small or large depending on whether environmental changes are small or large, respectively. Under an extreme environmental stress, the response may be also extreme and the phenotype may deviate far beyond the normal range. Examples of abnormal phenotypic changes during ontogenesis may be seen in development of diseases and malformation. An example that is easy to understand is habitual smoking that affects the DNA methylation states of the smoker (Breitling et al 2011; Alegria-Torres et al 2011). In addition, epigenetic causes have been suggested for type-2 diabetes, atherosclerotic heart diseases and various kinds of cancers (Lindblom et al 2015; Sandoval and Esteller 2012; Simo-Riudalbas and Esteller 2014). However, malformation and diseases are not directly related to evolution, and we will therefore not discuss these problems further.

Phenotypic changes that can lead to evolution will be those triggered by a large environmental change beyond the normal reaction norm. Such phenomena as “novelty” (Moczek 2008) or “innovation” (Muller and Newman 2005; Wagner 2012; Lickliter 2014) are concrete examples of extreme phenotypic changes. However, there are few observations of abnormal phenotypic changes that lead evolution. This is perhaps easily understandable. That is, if such a phenotypic change occurs adaptively, then it will immediately trigger evolution which completes in a relatively short period of time (the point (c) above; also see below); once the evolution completes, the new phenotype will have been genetically fixed, and hence becomes “normal” (and we cannot observe that phenotype as “abnormal”).

One of the few observed examples of significant phenotypic response to environment may be the Dutch Hunger Winter (Lumey et al 2007). The Hunger (1944-1945) brought an abnormal shortage of food and nutrition. Among the people who suffered from the Hunger, the most seriously affected were the fetus in early gestation. Cohort studies of the Dutch Hunger Winter revealed that many of these children, after becoming adults, developed life-style diseases such as obesity, heart diseases, and cancers (Roseboom et al 2006). These people also had specific epigenetic changes (such as DNA methylations) (Heijmans et al 2008), suggesting that epigenetic changes introduced in early gestation have persisted after birth until the development of diseases in adulthood. These epigenetic and phenotypic changes due to the environment (i.e., the Hunger) are not genetic, and thus, correspond to the F pathway in Fig. 1.

From an evolutionary view point, the issue is whether the epigenetic and phenotypic changes were adaptive to the environment. The historical fact that the people who were affected by the Hunger during early gestation developed various life-style diseases suggests that the answer is negative (Roseboom et al 2011). However, if we imagine the Hunger persisted indefinitely, then those affected people, being able to take up nutrition more efficiently than normal people due to the epigenetic and phenotypic changes, would have better survived. Thus, under the condition of extreme food shortage, the epigenetic phenotypic changes of those people might have been “adaptive.”

## 4 Evolutionary models and simulations

According to Fig. 1, the phenotype of an individual is realized through the genetic (G) and epigenetic/non-genetic (F) pathways. The phenotype of an individual is subject to natural selection (the arrow of “natural selection” in Fig. 1). It should be noted that when a particular individual is naturally selected during evolution its adaptive phenotype may be caused by the G pathway or the F pathway, or the combination thereof. But when the evolution is complete, the adaptive phenotype of all individuals in the population should be caused by the G pathway (the fixation of the adaptive genotype).

Based on the scheme represented in Fig. 1 we have proposed the “cooperative model” (Nishikawa and Kinjo 2014). The basic equation of this model is given as

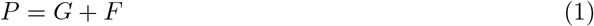

where the terms *P*, *G* and *F* represent the phenotypic, genotypic and epigenotypic values, respectively. The *G* and *F* terms correspond to the G and F pathways, respectively, in Fig. 1.

Following the standard treatment in quantitative genetics, if we assume that *P* represents the phenotypic value of a multifactorial trait, *G* is given as the sum of the contributions from multiple (possibly many) genetic mutations (Nishikawa and Kinjo 2014). It is assumed that *a significant amount of genetic mutations preexist in the gene pool of the population* (West-Eberhard 2005a). These mutations are assumed to be neutral before the environmental change after which a half of the mutations become advantageous and the other half disadvantageous. However, not all of the potentially advantageous mutations need to exist in any single individuals. In fact, individual minor mutations are generally scattered over different individuals and loci in the first generation, which is statistically more plausible. In other words, there may be no heritable adaptive trait in the first generation.

The *F* value of each individual is not inherited to its progeny, but in each generation it is assigned to each individual according to the same probability distribution (a Gaussian with a large standard deviation σ(*F*) and the zero mean) as long as the changed environment persists.

In comparison, the conventional model of Evolutionary Synthesis lacks the F pathway and considers (mostly) only the G pathway. The basic equation of the Evolutionary Synthesis (Barton et al 2007) is given by

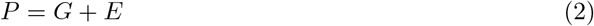

where *P* and *G* are common to Eq. 1, and *E* is the environmental deviation (also called the error term) given as a Gaussian variable with a small standard deviation σ(*E*) and the zero mean.

The only apparent difference between Eqs. (1) and (2) is the magnitude of standard deviations:

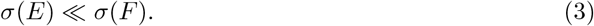

That is, the standard deviation of the *F* term is far greater than that of the *E* term.

The fundamental assumption of the Evolutionary Synthesis is that only heritable traits are subject to natural selection. As such, it has thoroughly avoided the contribution of acquired traits to evolution. However, it is the phenotype upon which natural selection operates, and the phenotype depends on both genome and epigenome (Fig. 1). In other words, there is no means to distinguish, at the phenotype level, whether a given trait is heritable or acquired. Therefore, it is more reasonable to conclude that an acquired trait, or the *F* term in Eq. (1), can also contribute to evolution.

Eq. (2) should be regarded as the basic equation of genetics rather than that of evolution. This equation is often used for studying genetic and environmental contributions to disease phenotypes (i.e., heritability) (Barton et al 2007). In the context of evolution, the Evolutionary Synthesis adopted this equation only formally, and the contribution of the *E* term is often assumed to be so small that it is completely ignored by setting *E* = 0 (Crow and Kimura 1970). Then, Eq. (2) reduces to *P* = *G* so that all the factors contributing to evolution are genetic and non-heritable acquired traits are completely excluded.

We compared the two models given by Eqs. (1) and (2) using computer simulations (Nishikawa and Kinjo 2014). We assumed that evolution started with a large environmental change, and studied the simulation trajectories of populations encountered with the persisting environmental change. The environmental change was assumed to have occurred at some point in time, before which a certain stable environment was assumed to have continued for a sufficiently long time, and after which the changed environment persisted indefinitely. The changed environment imposes a certain evolutionary pressure. Individuals with an adaptive phenotype overcome the pressure and survive, and those without are mostly eliminated from the population. In other words, we assumed natural selection by a threshold model. For the details of the simulations, refer to Nishikawa and Kinjo (2014).

The simulation results based on the cooperative model and the conventional model are summarized in Fig. 2. In the case of the conventional model (Eq. 2), the population size monotonically decreased and became extinct within a relatively small number of generations (Fig. 2A). On the contrary, in the case of the cooperative model, the population size decreased for the first several generations, but it recovered rather quickly to the original size (Fig. 2A). This result shows that the apparently small difference in the models (Eq. 3) leads to a great qualitative difference in the evolutionary trajectories.

**Fig. 2.**
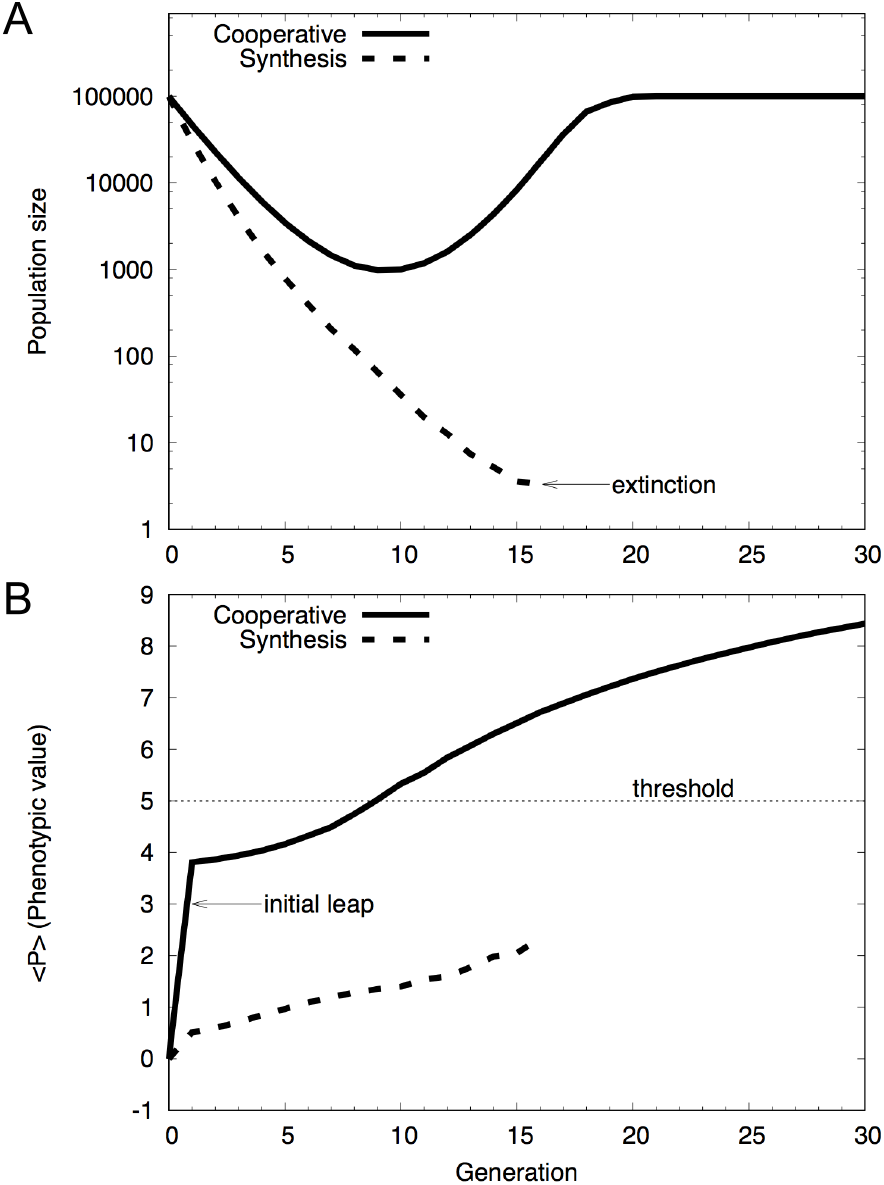
Simulation results. (A) Population changes caused by the environmental change occurred at the 0-th generation are plotted against the generation. (B) The average phenotypic value per individual, 〈*P*〉, is plotted against the generation. The solid curve is based on the cooperative model (Eq. 1), while the dashed curve on the Evolutionary Synthesis (Eq.2). The simulation has been performed using the genetic algorithm, where the genome of an individual, inherited from the parents, is randomly shuffled and those individuals with the phenotypic value higher than the threshold (*P* > *T*) are survived via the natural selection. The initial leap appeared in the cooperative model (B) is caused by the epigenetic plasticity of the organisms (see the text). Reproduced after Figure 1 of Nishikawa and Kinjo (2014) with modifications.

## 5 Initial leap/spread of phenotypic change

What is particularly noticeable in the simulations of the cooperative model is that the average phenotypic value per individual, 〈*P*〉, increases immediately in the first generation (Fig. 2B). This increase, which we refer to as the “initial leap”, reflects the immediate response of individuals to the environmental change without a generational change, and it is not accompanied with genetic mutations and therefore is purely of an acquired phenotypic change. West-Eberhard called it the “initial spread” (West-Eberhard 2005b), and stressed its importance in the initiation of a phenotype-driven evolution. After the initial leap, 〈*P*〉 monotonically increases as generations change.

The Evolutionary Synthesis claims that, initially, an individual with advantageous genetic mutations appear in the population, and through generational changes the advantageous individuals are selected to eventually fix the genetic mutations in the population. However, the first genetic mutation must appear in the germ line of a single individual in the population. Even if that mutation is advantageous, it seems extremely difficult for the mutation to spread over the entire population. Furthermore, if many genetic mutations are required to express an adaptive phenotype, the probability for an individual to have all the necessary mutations is extremely low. On the other hand, if a significant fraction of individuals in the population can express an advantageous phenotype at once without genetic mutations, just as in the case of the initial leap/spread, they are more likely to survive and to reproduce (Gilbert and Epel 2009). Thus, evolution can initiate without difficulty, contrary to the case of the Evolutionary Synthesis. In our simulation, 5% of the population exhibited the phenotypic values greater than the threshold (Nishikawa and Kinjo 2014). The precise value depends on the simulation parameters, but it is important that a non-negligible number of individuals were able to survive owing to an adaptive phenotype caused by epigenetic plasticity.

The emergence of phenotypic changes in many individuals of the very first generation is also seen in Waddington’s experiments of genetic assimilation. When a heat shock was applied to pupae, 40% of the population, when becoming adult, exhibited the *cross-veinless* phenotype (Waddington 1953). When ether was applied to fertilized eggs, 25%-50% of the surviving adults exhibited the *bithorax* phenotype (Waddington 1956). Such large fractions of phenotypic variants in the first generation suggest that they are induced by some epigenetic causes rather than cryptic genetic mutations (see below).

## 6 Mechanism of genetic assimilation

The simulations based on the cooperative model reproduce genetic assimilation. To see this more clearly, let us decompose the average phenotypic value 〈*P*〉 (Fig. 2) into the genetic (〈*G*〉) and epigenetic (〈*F*〉) components (Fig. 3). The average *F* value, 〈*F*〉, sharply increases in the first generation after which it gradually decreases. This shows that the epigenetic factor is indeed responsible for the initial leap of the phenotypic value (Fig. 2). On the contrary, the average *G* value, 〈*G〉,* is nearly 0 in the first generation, and increases monotonically and gradually in later generations. Finally, the *G* value by itself exceeds the threshold value, which indicates that the phenotypic change has been genetically assimilated. It is interesting to note that 〈*G*〉 constantly increases even during the period when the population size decreases (c.f., Fig. 2).

**Fig. 3.**
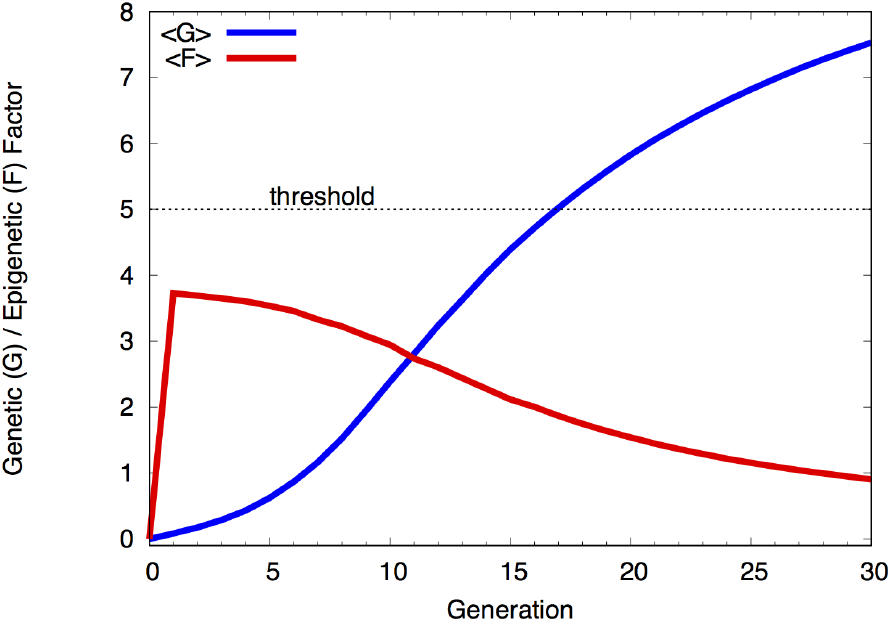
The average genotypic value per individual, 〈*G*〉, and the average epigenotypic value per individual, 〈*F*〉, plotted against the generation for the cooperative model (Eq. 1). Note that the initial leap and then gradual decrease of 〈*F*〉 is in contrast with the sigmoidal increase of 〈*G*〉. Recast the same data as those in Figure 1 of Nishikawa and Kinjo (2014).

The *G* value of individuals may increase or decrease relative to their parents as a result of genome shuffling during a generational change (Nishikawa and Kinjo 2014). However, genome shuffling alone neither increases nor decreases the average 〈*G*〉 value of the population as it is a random process with no preference. For the 〈*G*〉 value to increase, natural selection must act on the phenotype which consists of both *G* and *F*. In the first several generations, the *G* value alone is not large enough to overcome the selection threshold on average and a large *F* value is necessary for an individual to survive (Fig. 2). As a result of genome shuffling, some individuals may have relatively larger *G* values. Since the *F* value is assigned to all individuals according to the same probability distribution, those with higher *G* values are on average more likely to be naturally selected. This also implies that as the *G* value increases, a smaller *F* value becomes sufficient to overcome the threshold (Eq. 1). Thus, advantageous genetic mutations increasingly accumulate via generational changes in single individuals (increase in 〈*G*〉) through the action of natural selection, and consequently, the epigenetic contribution to the phenotype decreases on average. We believe that this behavior of the cooperative model explains the mechanism of genetic assimilation.

What is important for genetic assimilation to occur is the “equivalence” of the G and F pathways with respect to the phenotype (Fig. 1) in that changes in either *G* or *F* may express the same phenotype. Although indistinguishable at the phenotype level, these phenotypic changes are clearly different at the molecular level. Genetic assimilation is a process in which the average epigenetic phenotypic change, 〈*F*〉, is replaced with the average genetic phenotypic change, 〈*G*〉, through generations subject to natural selection. Thus, the central problem West-Eberhard left regarding the mechanism of genetic assimilation has been solved.

## 7 Mutual independence between the G and F pathways

In addition to the equivalence of the G and F pathways, another crucial assumption is that the two pathways are mutually independent. Of course, the F pathway does depend on the genome in the sense that epigenetic machineries such as chromatin modification enzymes and non-coding RNAs are encoded in the genome. Similarly, the G pathway depends on epigenomes in the sense that gene expression is modulated by the epigenome. By the mutual independence, we mean that the F pathway (phenotypic response to the environmental change) does not depend on those genetic mutations that induce an adaptive phenotype (G pathway) or *vice versa.* We elaborate this point in the following by comparing the models studied by Lande (2009).

Lande (2009) proposed a model for evolution by genetic assimilation in which the phenotypic plasticity is caused by the reaction norm (adaptive response to an environmental change) and the reaction norm itself can evolve (thus depends on genetic mutations). He compared this model with two other models: a purely Darwinian model which is mostly equivalent to the one defined by Eq. 2, and the “Baldwin effect” model in which the reaction norm does not evolve but it causes a plastic phenotypic response that is always adaptive and constant (Price et al 2003).

For the Darwinian model, the phenotypic value evolves slowly after the large environmental change and the adaptive phenotype is eventually fixed in the population the size of which is assumed to be infinite. However, the fitness drops sharply and greatly at the environmental change so that the population of a finite size would become extinct as in our conventional model (Eq. 2).

For the Baldwin effect model, the (adaptive) phenotypic plasticity due to the reaction norm causes what is equivalent to the initial leap of the phenotypic value, which helps the population to survive. However, even when the advantageous genetic mutations are fixed in the population, the plastic response to the changed environment remains. In this sense, the adaptive phenotype is not genetically assimilated in this model.

In order to overcome this difficulty with the Baldwin effect model, Lande (2009) assumed that the (slope of) reaction norm could also evolve. Lande’s model did reproduce genetic assimilation in a sense: after the initial leap, the phenotypic value continued to increase due to the genetic mutations in the reaction norm (Phase 1), and in later generations, the plastic response decreased and replaced by non-plastic adaptive phenotype (Phase 2). However, it took ∼ 10^3^ generations before the Phase 2 started and ∼ 10^4^ generations was required for the genetic assimilation to complete, which is far longer than Waddington’s experiments (∼ 20 generations) or the cooperative model (10^1^ ∼ 10^2^ generations). It appears that the extensive adaptation by the evolution of reaction norm slowed down the evolution of non-plastic adaptive phenotype.

It should be pointed out that the Baldwin effect model is a special case of Lande’s model where the genes responsible for the reaction norm happen to be immutable. But the plastic response being always adaptive implies that the genome “expects” the changed environment (i.e., the response is genetically programmed). In this sense, the phenotypic plasticity of the Baldwin effect model should correspond to the G pathway (starting from the blue environmental cue in Fig. 1) rather than the F pathway. By the same token, Lande’s model deals with the evolution of the G pathway only. Conversely, if we assume the F pathway is independent of the G pathway, then the phenotypic response due to the former should not be always adaptive but *ad hoc* or stochastic, as assumed in the cooperative model. Because *F* is stochastic (with a large variance), the gradual increase in the average genotypic value 〈*G*〉 consequently makes the *average* epigenotypic value 〈*F*〉 decrease, which allows rapid evolution by genetic assimilation.

## 8 Cryptic variants vs. epigenetic changes

It has been suggested that Waddington’s experiments may be reproduced by experiments using Hsp90 mutants. In the following, we compare the two experiments and examine if the Hsp90 experiment really accounts for the molecular basis of Waddington’s genetic assimilation (Pigliucci 2003; Crispo 2007).

When heat shock is exerted on fruit flies, Hsp90 (a heat-shock protein) is over-expressed, which stabilizes various proteins that are destabilized by the heat shock and assist the folding of newly synthesized proteins. In the absence of heat shock, Hsp90’s assist the folding of mutant proteins and keep them functionally stable. Thus, even though some proteins contain minor mutations, their effect remains latent because of the function of Hsp90, and hence they are called “cryptic variants” (Gibson and Dworkin 2004).

Rutherford and Lindquist (1998) used in their experiments fruit flies with compromised Hsp90 such as those with Hsp90 mutant or the wildtype fed on a Hsp90 inhibitor (geldanamycin). When the flies with mutant Hsp90 are bred with various strains of flies to produce the F1 generation, a small fraction of this generation contained various types of phenotypic variants. This is understood as the expression of cryptic variants due to the compromised function of Hsp90 as the capacitor (Rutherford and Lindquist 1998). The types of cryptic variants are determined by the wildtype strains of breeding partners, and consequently, the phenotypic variants in the F1 generation are also determined by the strains. When a certain phenotypic variant is artificially selected and bred so that the polygenes responsible for the phenotype are enriched, after several generations, even those flies with normal Hsp90 exhibited the phenotypic change. This series of experiments may suggest that the initial phenotypic change has been genetically fixed after the cycles of artificial selection and generational changes.

We argue in the following that the Hsp90 mutant experiment is not an example of genetic assimilation. In short, the phenotypic variants induced by the compromised Hsp90 are all genetic mutants, and there seems to be no epigenetic factors involved in these experiments.

Firstly, the fact that different mutants were derived from different strains (Rutherford and Lindquist 1998) indicates that different strains contain different cryptic variants. This suggests that the phenotypic variants are due to not epigenetic, but genetic causes.

Secondly, in the Hsp90 experiment, only 2% of the first generation after the breeding between Hsp90-mutant and wildtype showed phenotypic changes (Rutherford and Lindquist 1998), which is reasonable if we assume a polygenic phenotypic change (Rutherford and Lindquist 1998). In Waddington’s experiments, on the contrary, a relatively large fraction of the population exhibited the phenotypic change in the first (F0) generation of the artificial selection (e.g., in the case of the *bithorax* experiment, the fraction of mutants was 25%-50% depending on strains), suggesting that the *bithorax* mutants are caused by epigenetic mechanism. In addition, the memory effect (Cheedipudi et al 2014; Vickers 2014) that the exposure of eggs to ether caused the *bithorax* mutation in adulthood suggests an epigenetic origin of the mutant.

Finally, in the Hsp90 experiment, only 5 or 6 generations were required for the mutant to be fixed in the population (Rutherford and Lindquist 1998), which is evidently less than that in Waddington’s experiment (13-20 generations). In the former, since artificially selected mutants always include minor genetic mutations contributing to the mutant phenotype, the repeated application of artificial selection only to such mutants is expected to rapidly enrich the polygenes. On the contrary, in Waddington’s experiments, artificially selected phenotypic variants do not necessarily have genetic mutations. Particularly in the first several generations, epigenetic contribution to the phenotype is more prominent than genetic one while the genetic contribution increases only more slowly (Fig. 3) compared to the artificial selection of purely genetic mutants.

Apart from the Hsp90-as-a-capacitor model, it has been reported that Hsp90 is primarily involved in an epigenetic regulatory system so that compromised Hsp90 alters chromatin regulation (Sollars et al 2003; Sawarkar and Paro 2013). In fact, mutants caused by the loss of function of Hsp90 include various types of phenotypic abnormalities. Therefore, it is plausible that some phenotypic variants are due to genetic mutations and others are due to epigenetic factors. In the latter case, the mechanism suggested by the cooperative model should apply.

There are many observations that may be considered as concrete examples of naturally occurring genetic assimilation (Schlichting and Wund 2014; Matsuda 1987). It seems difficult to explain all these examples solely in terms of the Hsp90-as-a-capacitor model. Genetic assimilation should be considered as a far-reaching concept that is applicable to phenotype-driven and adaptive evolution in general. At the same time, it is a general evolutionary mechanism that incorporates epigenetic factors as an essential element in addition to genetic mutations.

## 9 Conclusion

Epigenetics, a rapidly evolving field in biology, has enabled us to examine the genotype-phenotype relationship from a whole new perspective. According to epigenetics, the phenotype of an organism depends more strongly and directly on epigenome than genome or genotype. This fact is plainly evident in the relationship between cell differentiation and epigenome in multicellular organisms. Environment can affect and alter epigenomes, and the altered epigenomes are conserved through cell divisions and their effect persist throughout the life time of an organism. Conventional Evo-Devo theories have well recognized the importance of phenotypic plasticity. Now, they may be recast as a molecular mechanism connecting from environment to epigenome to phenotype. Thus, epigenetics is closely linked to evolution through phenotypic plasticity. The phenotype of an individual, which is subject to natural selection, arises from a combination of genetic (*G)* and epigenetic (*F)* factors. The simulations based on such a mechanism, i.e., the cooperative model, reproduced the process of genetic assimilation. The results of the simulations have suggested that genetic assimilation is a process of generational changes in which the average 〈*F*〉 per individual in the population is being replaced with the average 〈*G*〉 per individual by the action of natural selection. Finally, it is noted that the epigenetic changes causing the phenotypic change in the first generation occur in somatic cells. They cannot therefore be transferred to the next generation by transgenerational epigenetic inheritance (although TGEI itself is not inconsistent with genetic assimilation).

## Compliance with ethical standards

### Ethical approval

This article does not contain any studies with human participants or animals performed by any of the authors.

